# Hippocampal microcircuits constrain the generation of epileptiform activity

**DOI:** 10.64898/2026.06.25.734494

**Authors:** Quynh-Anh Nguyen, Jordan S. Farrell, Barna Dudok, Gergely G. Szabo, Tilo Gschwind, Jesslyn Homidan, Ivan Soltesz

## Abstract

Epilepsy is largely characterized using macroscale measures of neural activity, such as electroencephalography, which are unable to resolve the underlying cellular-level substrates of pathological activity. Although there is a growing understanding that hippocampal microcircuits, comprised of distinct CA1 principal cells (PCs), inhibitory neurons, and input/output relationships, route information through parallel hippocampal pathways, the relevance of this microscale organization in epilepsy is poorly understood. To address this gap, we focally knocked out the β3 subunit of the GABA_A_ receptor from CA1 PCs, which strongly impaired parvalbumin-mediated inhibition to deep PCs, but not superficial PCs – potentially promoting a microcircuit-selective epilepsy manifestation. Indeed, we observed robust interictal epileptiform discharges (IEDs) at the site of focal knockout with features consistent with a potential CA2-to-deep CA1 PC generation mechanism. In line with prior observations that CA2 inputs potently drive deep PCs under physiological conditions, IEDs could be reliably evoked by optogenetic stimulation of CA2 in β3 focal knockout mice. Resolving cellular activity across the CA1 PC network during IEDs, *in vivo* 2-photon calcium imaging demonstrated stronger activation of deep versus superficial PCs. Altogether, these data demonstrate that macroscopic electrophysiological patterns such as IEDs can have underlying microcircuit constraints, which has important implications for designing therapies that target the underlying microcircuit generators of epileptiform activity.

## Introduction

Conventional methods to measure epileptiform activity record at a scale far removed from the circuit mechanisms that are thought to generate pathological activity patterns^1–3^. Resolving the underlying generators of epileptiform activity at the appropriate scale could inform why some patients with ostensibly similar epilepsies respond to medications and others do not. The hippocampus and temporal lobe structures, for instance, are associated with especially high rates of drug resistance and widely implicated in adult focal epilepsy^4,5^. Several studies under physiological conditions have brought forth the concept that the hippocampus is composed of microcircuits that give rise to parallel pathways for information routing^6–8^. This is most appreciated in the CA1, where deep and superficial principal cells (PCs) have distinct developmental origins, molecular markers, excitatory inputs, inhibitory motifs, information encoding, and projection targets^7–16^. Whether these CA1 microcircuits can function as distinct generators of epileptiform activity is unknown but is crucial to our understanding of epilepsy mechanisms.

To address this gap in understanding between hippocampal microcircuits and macroscale electrographic patterns, we leveraged the fact that inhibition onto deep and superficial PCs are remarkably divergent^10^, and that loss of inhibition is a common driver of epilepsy^17–21^. Previous work found that parvalbumin (PV)-expressing basket cells (BCs) strongly inhibit deep CA1 PCs and only weakly inhibit superficial PCs^10^. Since PVBCs are potent controllers of seizure activity^22^ and thought to be a key chokepoint in epilepsy^3,23^, we hypothesized that loss of this normally strong inhibition onto deep CA1 PCs would result in epileptiform activity emerging primarily from this microcircuit. Since the β3 subunit of the GABA_A_ receptor is a main contributor of PV-mediated inhibition in CA1 PCs^24^ and mutations in the gene encoding this subunit, *GABRB3*, can drive severe epilepsy syndromes^25^, we employed a focal knockout strategy to impair PV-mediated inhibition onto deep CA1 PCs. Here we demonstrate that focal loss of *GABRB3* in CA1 PCs selectively impairs PV-mediated inhibition onto deep PCs and results in local interictal epileptiform discharges (IEDs). We also provide evidence that IEDs are associated with stronger activation of deep CA1 PCs than superficial PCs and can be reliably evoked by stimulating canonical excitatory afferents onto deep CA1 PCs. Collectively, this work demonstrates that IEDs can be constrained by microcircuit-level organization, which may regulate differential manifestations of epilepsy in otherwise similar focal epilepsies.

## Results

To perform a focal knockout of the β3 subunit of the GABA_A_ receptor in CA1 PCs, referred to here as β3 fKO, we injected AAV-CaMKIIa-mCherry-Cre virus (or AAV-CaMKIIa-mCherry as a control) into the CA1 of adult *GABRB3*^*fl/fl*^ mice^26^ (**Figure 1A**). We first confirmed that Cre was reliably transduced by examining mCherry reporter expression and that β3 expression, measured by immunofluorescence, was abolished in the CA1 at the injection site (**Figure S1**). Next, we tested the hypothesis that PV-mediated inhibition onto deep PCs would be selectively compromised in β3 fKO mice. To this end, we performed whole-cell patch clamp recordings of deep and superficial CA1 PCs in double transgenic β3^fl/fl^/PV^Flp^ mice injected as above alongside Ef1a-fDIO-ChR2-eYFP (**Figure 1A**). Optogenetically-elicited PV-mediated inhibitory currents were larger in deep versus superficial CA1 PCs in control mice, reflecting the cell-type specific connectivity previously found^10^ (**Figure 1B,C**). In contrast, deep CA1 PCs from β3 fKO mice had virtually no response to PV cell activation (**Figure 1B,C**). Importantly, superficial PCs recorded from the same β3 fKO slices maintained optogenetically-evoked inhibitory currents (**Figure 1B,C**), demonstrating that the impaired PV-mediated inhibition preferentially affects deep PCs and is not related to differences in the optogenetic or slice preparation. These results demonstrate that focal loss of *GABRB3* in CA1 robustly reduces the normally strong PV-mediated inhibition onto deep PCs.

**Figure 1:**
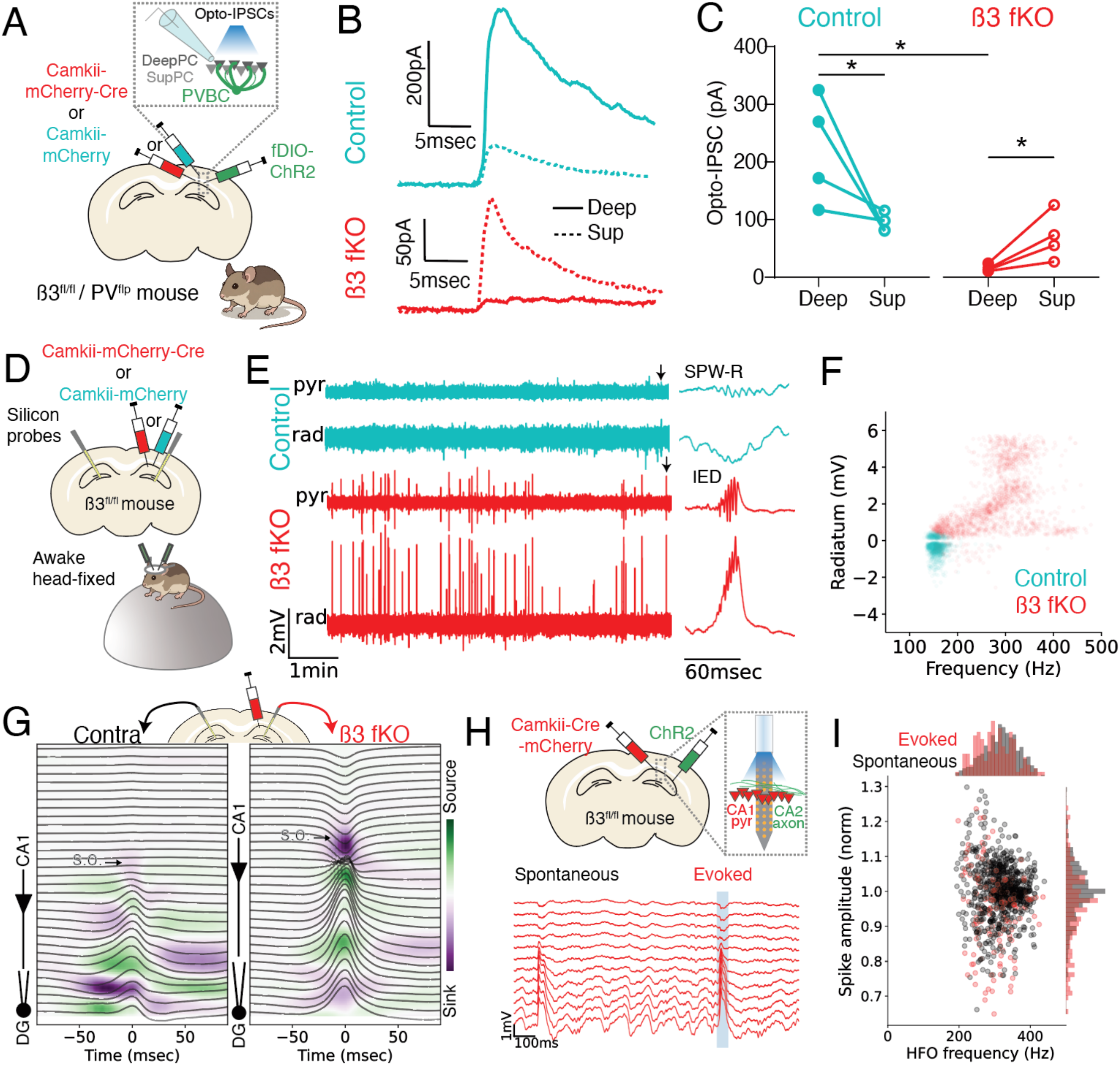
Impaired inhibition onto deep CA1 PCs and emergence of IEDs in β3 fKO mice. (A) β3^fl/fl^/PV^Flp^ mice were injected with AAVs to knockout β3 (or control) and express ChR2 in PV cells. Whole-cell recordings were done in both deep and superficial cells within the same slice and PV-mediated IPSC responses were elicited by blue light stimulation. (B) Representative traces of whole cell recordings in deep and superficial (Sup) cells in control and β3fKO mice. (C) Quantification of B. Deep cells from control mice receive stronger PV input than superficial cells (t3=2.383, p=0.0487), whereas superficial cells receive stronger input than deep cells in β3 fKO mice (t3=3.044, p=0.0278). This difference is also seen between subjects, with deep cells in β3 fKO mice showing decreased PV input compared to deep cells in control mice (t6=4.361, p=0.0219,) while responses in superficial cells are not significantly different between controls and β3 fKOs (t6=1.056, p=0.3517). (D) β3 fKO and control mice were recorded with 32-channel silicon probes from both hippocampi while awake, head-fixed, and free to move on floating styrofoam ball. (E) Representative LFP traces from the stratum pyramidale (pyr) and radiatum (rad) from control and β3 fKO mice. Arrows denote a representative SPW-R and IED, shown on the right on an expanded timescale. (F) Summary of detected events in the ripple and HFO bands (100-500Hz), showing mean oscillatory frequency and maximal radiatum voltage for control (n=8 mice, 1378 events) and β3 fKO mice (n=11 mice, 1964 events). Oscillatory frequency: t17=5.93, p=0.0000163. Radiatum voltage: t17=4.12, p=0.000714. (G) Average current source density and LFP aligned to the peak of IEDs in a representative β3 fKO mice. In the CA1, dominant sinks (purple) are observed in stratum oriens where CA2 axons terminate. s.o. = stratum oriens (H) β3 fKO mice were injected with AAV-ChR2 in CA2. Blue light pulses delivered to CA2 axons at the CA1 recording site evoked IEDs with similar properties to spontaneous IEDs. (I) Quantification of H for HFO frequency and spike amplitude for spontaneous (black) and optogenetically elicited IEDs (red). n=3 mice, p=0.08 (frequency) and p=0.22 (amplitude).

Since loss-of-function variants in *GABRB3* can cause epilepsy^25^, we next ascertained whether focally-restricted β3 loss in CA1 would result in pathological activity patterns *in vivo*. Bilateral silicon probe recordings were performed on awake head-fixed β3 fKO and control mice at least two-weeks post-AAV injection (**Figure 1D**). We observed frequent IEDs only in β3 fKO mice, composed of a large positive spike in the stratum radiatum, unlike a typical negative sharp-wave during physiological ripples in control mice^27^, and a high-frequency oscillation (HFO, >200Hz^28^) in the pyramidal cell layer (**Figure 1E, F**). HFOs were largely not observed in the contralateral hemisphere of β3 fKO mice, confirming their focal manifestation in the AAV-transduced region (**Figure S2**). Despite the presence of clear epileptiform activity, β3 fKO mice did not exhibit gross anatomical changes or neuronal loss, however increased microglial density was noted (**Figure S3**). Altogether, these data support that β3 fKO mice offer a unique model to study the role of selectively perturbed microcircuits without the confounds of hippocampal sclerosis seen in more commonly used rodent models of focal temporal lobe epilepsy (e.g., pilocarpine^29^ or intrahippocampal kainate^30^).

Under physiological conditions, SPW-Rs are generated by the highly recurrent and bilaterally projecting CA3 network, which produce ripple oscillations that are typically bilaterally synchronous^27,31^. Consistent with prior studies^31,32^, 64% of ripples were detected bilaterally in control mice (**Figure S2**). In β3 fKO mice, however, HFOs co-occurred with SPW-Rs in only 28% of cases (**Figure S2**), highlighting other potential circuit mechanisms that contribute to IED generation beyond the ripple-producing CA3 network. To understand the possible circuits involved in IED generation, we first performed current source density analysis on the laminar recordings across CA1 in β3 fKO mice. IEDs were associated with a prominent current sink in stratum oriens, which was also observed contralaterally (**Figure 1G**). This pattern is consistent with strong excitatory input to the basal dendrites of CA1 PCs, where CA2 axons terminate, rather than afferent input from CA3. Interestingly, prior work found that deep PCs receive roughly 3-fold stronger input from CA2 afferents than superficial PCs^33^. Along with the loss of PV-mediated inhibition to deep CA1PCs, these data point to CA2 as a potential driver of IEDs in β3 fKO mice. Indeed, optogenetic activation of CA2 axons reliably evoked IEDs in β3 fKO mice (**Figure 1H**). Importantly, the characteristics of evoked IEDs were indistinguishable from spontaneous IEDs, suggesting similar underlying mechanisms (**Figure 1I**).

To further resolve the recruitment of distinct CA1 PC microcircuits during IEDs in β3 fKO mice, we utilized *in vivo* 2-photon calcium imaging to record from large ensembles of CA1 PCs in awake mice while they were free to run or rest on a treadmill (**Figure 2A**). Compared to control mice, which generally had the highest calcium activity during running periods, neuronal activity in β3 fKO mice was greatest during brief events occurring while mice were immobile (**Figure 2B**). To capture these bursts of activity during immobility, hypersychronous events were defined as times in the recording when the percentage of active cells during immobile periods exceeded that of locomotion (see methods). While few events were detected in control mice, β3 fKO mice displayed a higher rate of hypersynchronous events and a greater proportion of active cells within these events (**Figure 2C,D**). Using simultaneous LFP in a subset of recordings, we confirmed that these brief periods of high network calcium activity are accompanied by an IED in β3 fKO mice (**Figure 2E**). We then plotted the average calcium activity during hypersynchronous events for each CA1 PC in a representative β3 fKO mouse, noting the location of stratum radiatum and oriens to determine the superficial versus deep axis (**Figure 2F**). Consistent with a hyperactive deep CA1 PC population, the most active cells during hypersynchronous events in β3 fKO mice were anatomically clustered towards the deep axis of this gradient (**Figure 2F**). By segregating CA1 PCs along this axis, we observed that deep CA1 PCs were more active than superficial PCs for all β3 fKO mice (**Figure 2G**). Thus, deep CA1 PCs are hyperactive during pathologically synchronous events, likely owing to reduced PV-mediated inhibition and strong drive from CA2.

**Figure 2:**
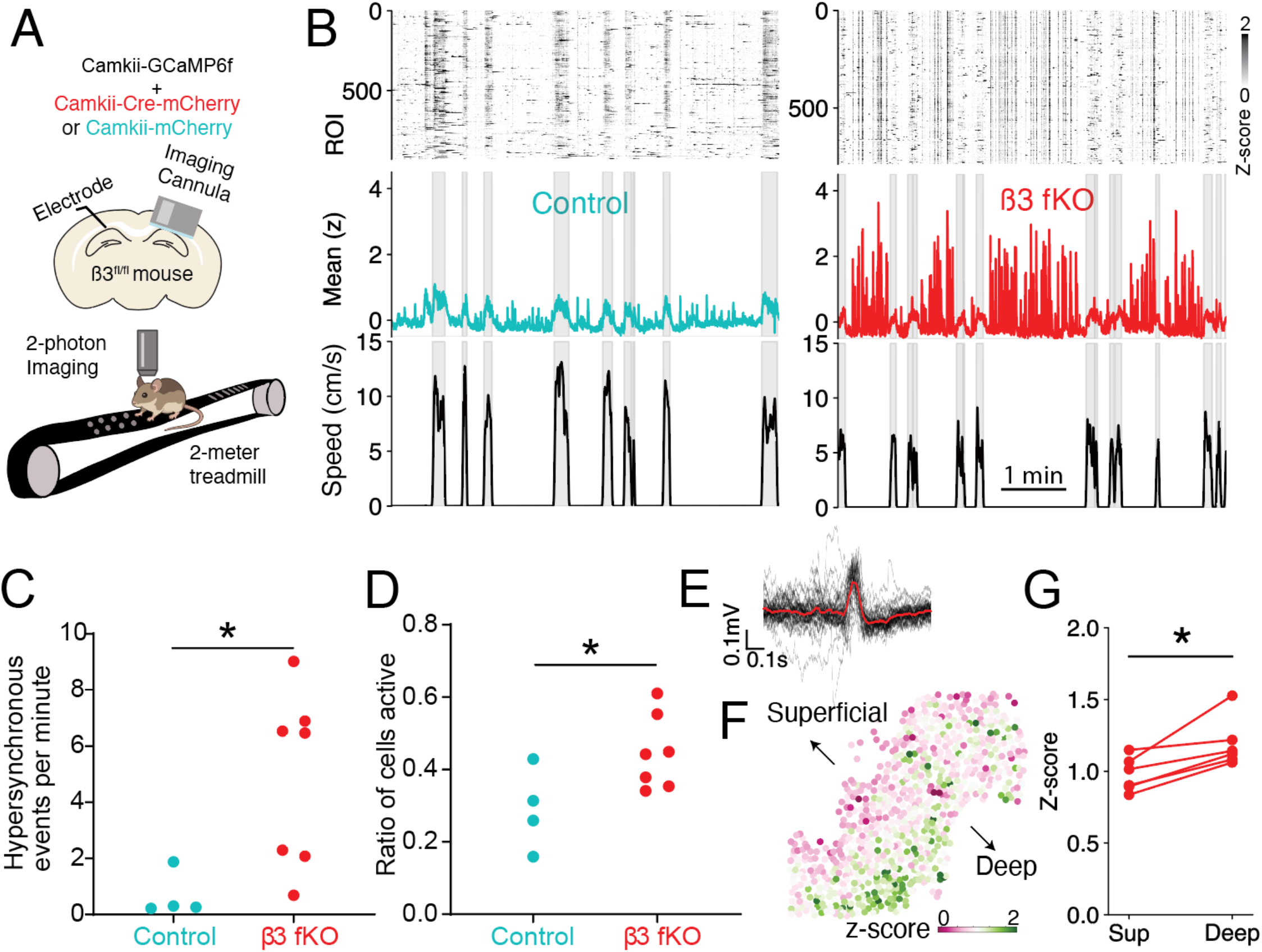
Synchronous events occurring during immobility are associated with strong activation of deep CA1 PCs in β3 fKO mice. (A) β3^fl/fl^ mice were injected with AAVs to knockout β3 (or control) and express GCaMP6f in CA1 PCs. 2-photon calcium and electrophysiological data were collected while mice ran or rested on a texture-cued treadmill. (B) Representative recordings from control and β3 fKO mice. Top shows individual traces, middle shows mean activity across cells, and bottom shows locomotor speed. (C) β3 fKO mice show an elevated frequency of large, synchronous calcium events during immobility (p=0.0121). (D) Higher ratio of active cells in β3 fKO mice versus control mice. (p=0.0424). (E) Simultaneous LFP was aligned to synchronous calcium events from a representative session in a β3 fKO mouse. Each black trace is LFP surrounding a single event, whereas red is the mean. (F) Calcium activity (z-score) shown as color for each cell during synchronous calcium events during immobility from a representative β3 fKO mouse. The deep vs. superficial axis is noted, with stronger recruitment observed in deep CA1 PCs. (G) Calcium activity (z-score) is greater for deep than superficial cells during synchronous calcium events during immobility. t5=4.74, p=0.0052.

## Discussion

Here we demonstrate that focal knockout of the β3 subunit of the GABA_A_ receptor compromises the normally strong PV-mediated inhibition onto deep CA1 PCs, resulting in microcircuit-selective hyperexcitability and IEDs emerging via the CA2-to-deep CA1 PC microcircuit. While this work encompasses one gene in one circuit, providing a level of control for mechanistic insights, the concept of microcircuit-selective epilepsy expression could more generally apply to other perturbations and other circuits. For example, with the advent of large-scale transcriptomic datasets^34–37^, it is now more feasible to assess how genes implicated in epilepsy are differentially expressed in microcircuits of interest. Given that microcircuits in CA1 are connected to vastly different large-scale networks^7^, expression differences along this axis of CA1 PC function could play an important role in the overall manifestation of seizures.

PVBCs are highly active during SPW-Rs, often reaching firing rates exceeding 100Hz^38,39^, which is thought to be crucial in regulating ripple oscillations^40^. Indeed, prior work found that HFOs can emerge within the same networks that drive SPW-Rs, but when inhibition is compromised, for example by GABA_A_ receptor blockade or when PVBCs enter depolarization block^40,41^. Unlike prior pharmacological approaches that broadly impair inhibition, our genetic perturbation has a strongly biased effect on PV-mediated inhibition onto deep CA1 PCs. Since deep CA1 PCs receive more PV-mediated inhibition^10^ and fire significantly less during SPW-Rs^12^, the observation that local IEDs emerge during contralaterally-recorded SPW-Rs underscores the important role of PVBCs in constraining activity during SPW-Rs. However, many IEDs occurred without a contralaterally detected SPW-R, which does not fit with the model that HFOs emerge from canonical CA3-driven SPW-R mechanisms. An alternative CA2-dependent mechanism may drive a subset of IEDs and is interesting to consider given the growing understanding of its role as a resilient seizure-generator in temporal lobe epilepsy^42–44^. Interestingly, CA2 PCs are highly active during immobility under normal conditions and have been implicated in a variety of synchronized hippocampal events, such as SPW-Rs with a stratum oriens sink^45^, an “n”-wave where non-ripple coupled cells predominantly fire^46^, and the recently described barrage of action potentials (“BARR”) generated by bursts of CA2 PCs^47^. Although more work is required to delineate how CA2-driven circuit patterns contribute to the manifestation of epileptiform activity, these data highlight the need to look beyond the conventional CA3 to CA1 circuit^48^.

Given the growing understanding of the genetic contributions to epilepsy and the expression of these genes within hippocampal microcircuits, this work provides a critical bridge in understanding by demonstrating how genetic perturbations in a given brain region can distinctly affect the microscale function of that circuit to generate macroscopic epileptiform patterns. Further work on understanding how microcircuit organization provides a conduit for epileptiform activity across various seizure models could help guide more precise therapies. Rather than gene therapy approaches for each genetic diagnosis, altering the excitability of select microcircuits may be a more general approach to treating epileptiform activity that converge on the same microcircuit. Directly related to this work and building from the growing understanding of the role of CA2 in acquired epilepsy models^42–44^, the CA2-to-deep CA1 PC microcircuit may be an important target for seizure therapy.

## Resource availability

### Lead contact

Requests for further information and resources should be directed to and will be fulfilled by the lead contacts, Quynh Anh Nguyen and Jordan Farrell.

### Materials availability

This study did not generate new unique reagents.

## Data and code availability

- All data reported will be shared by the lead contact upon request.
- Open-source code used is acknowledged in the methods
- Additional information and code required to reanalyze the data reported in this paper is available from the lead contact upon request.

## Acknowledgments

We thank Theresa Nguyen, Sylwia Felong, and Anna Ortiz for technical assistance with this work. This work was supported by the Esther A. & Joseph Klingenstein Fund (J.S.F. & Q.A.N.), Simons Foundation (J.S.F.), CureSHANK (J.S.F.), Citizens United for Research in Epilepsy (J.S.F.), McNair Medical Institute at The Robert and Janice McNair Foundation (B.D.), Whitehall Foundation (B.D.), and the National Institutes of Health (NIH) (grants NS131728, NS121106, NS136355 to I.S.; NS007280, NS106764, NS121399 to Q.A.N.; NS126725 to J.S.F.; and NS117795 to B.D.).

## Author contributions

Conceptualization: Q.A.N., J.S.F., and I.S.

Methodology: Q.A.N., J.S.F., B.D., G.G.S., T.G., and J.H.

Investigation: Q.A.N., J.S.F., B.D., G.G.S., and J.H.

Writing – Original Draft: Q.A.N., J.S.F.

Writing – Review & Editing: All authors

Supervision: I.S.

## Supplemental Information

**Figure S1:**
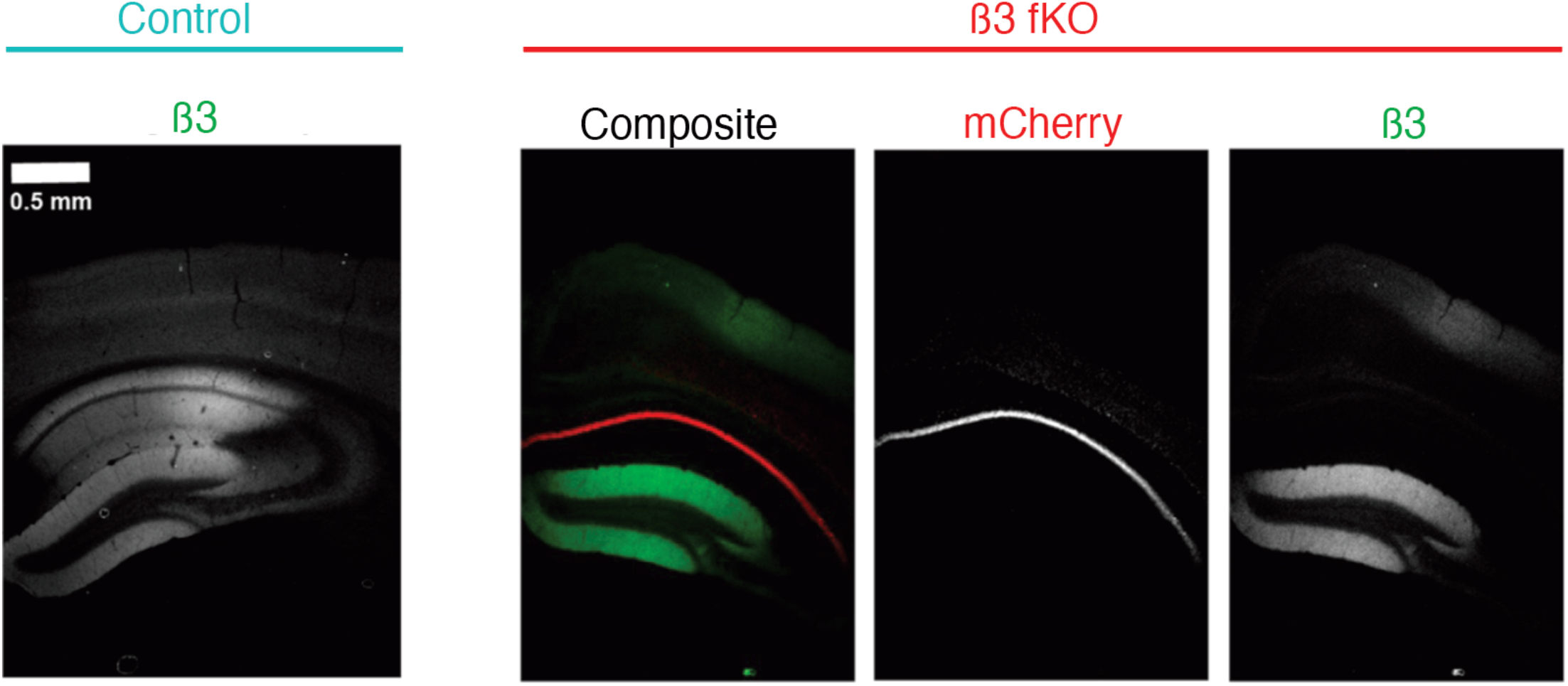
Focal loss of β3 in *Gabrb3fl/fl* mice following AAV-CaMKII-Cre-mCherry injection. Compared to control mice with abundant β3 expression, β3 fKO mice display an absence of β3 expression. Notably, β3 expression is spared in the dentate gyrus, confirming the specificity of the knockout strategy.

**Figure S2:**
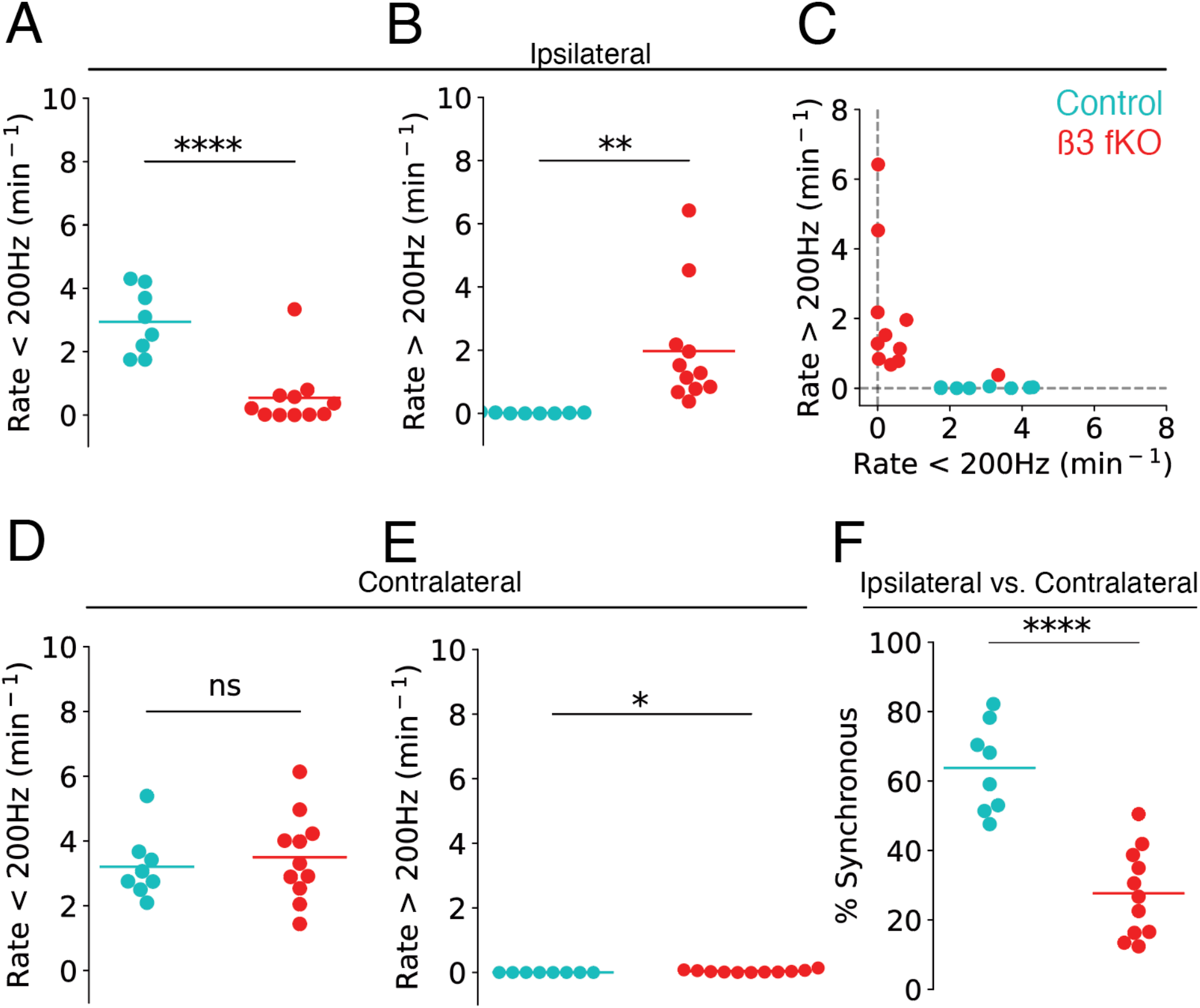
Pathological HFOs localize to the ipsilateral CA1 in β3 fKO mice. (A) β3 fKO mice have fewer physiological ripples (<200Hz) than control mice ipsilateral to the AAV injection site. t_17_ = 5.16, p = 0.0000791. (B) β3 fKO mice have more pathological HFOs (<200Hz) than control mice ipsilateral to the AAV injection site. t_17_ = 2.95, p = 0.00902. (C) Data from A and B replotted. An exponential function better fits the data than a linear function (r=0.72 vs. r=-0.60, respectively), suggesting that SPW-Rs and HFOs tend to not co-occur in the same region. (D) No differences were observed in the rate of contralateral physiological ripples. t_17_ = 0.51, p = 0.614. (E) β3 fKO mice have more pathological HFOs (<200Hz) than control mice on the contralateral hemisphere, but the average event rate is very low (0.04 ± 0.012 events per minute in β3 fKO mice versus 0.0 in control mice). t_17_ = 2.64, p = 0.0171. (F) Percent of events, inclusive of physiological ripples (<200Hz) and pathological (>200Hz) HFOs, that are coincident with a contralaterally-detected SPW-R. t_17_ =6.07, p = 0.0000126. *p<0.05, p<0.01, ****p<0.0001

**Figure S3:**
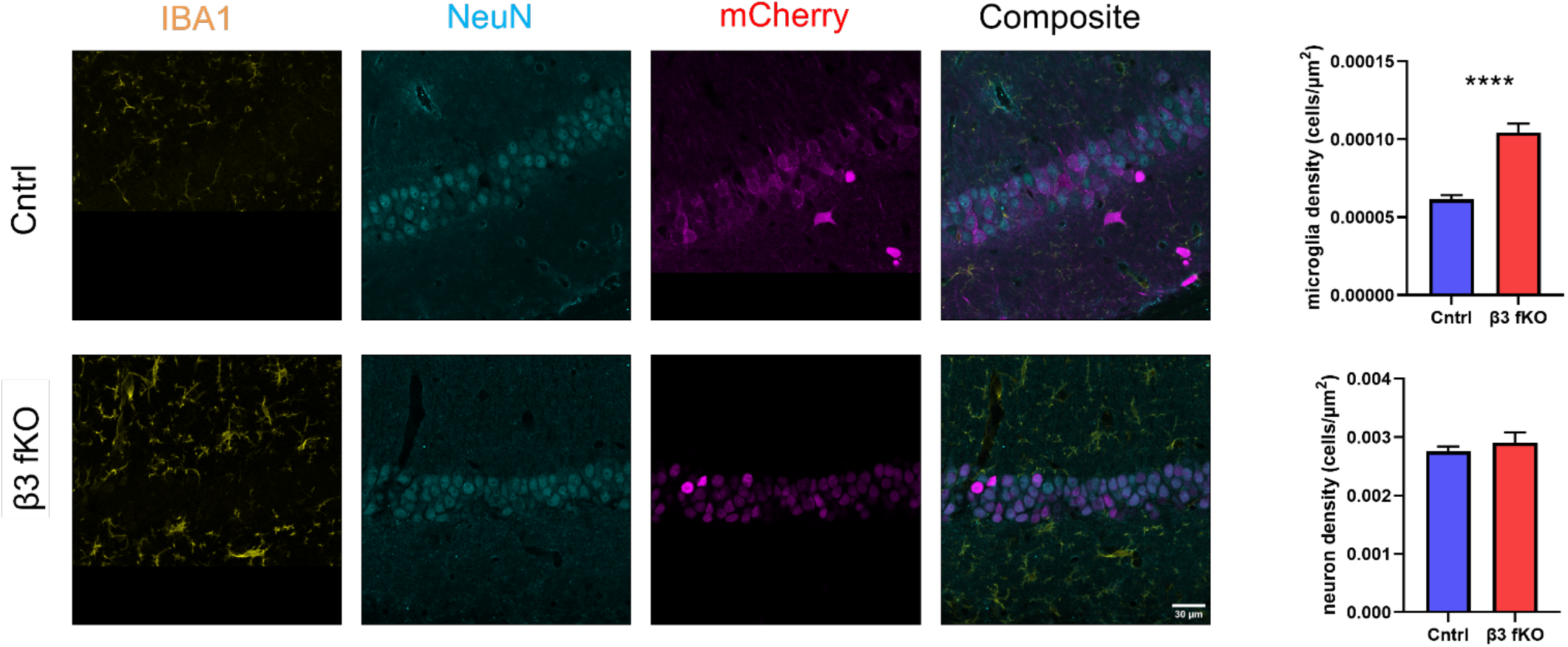
Histological characterization of inflammation and cell death in β3 fKO mice. Increased inflammation (p<0.0001) but no cell death (p=0.9742) observed in CA1 tissue from β3 focal knockout mice (n=12 sections/4mice control, 10 sections/4mice β3 focal knockout).

## Key resources table

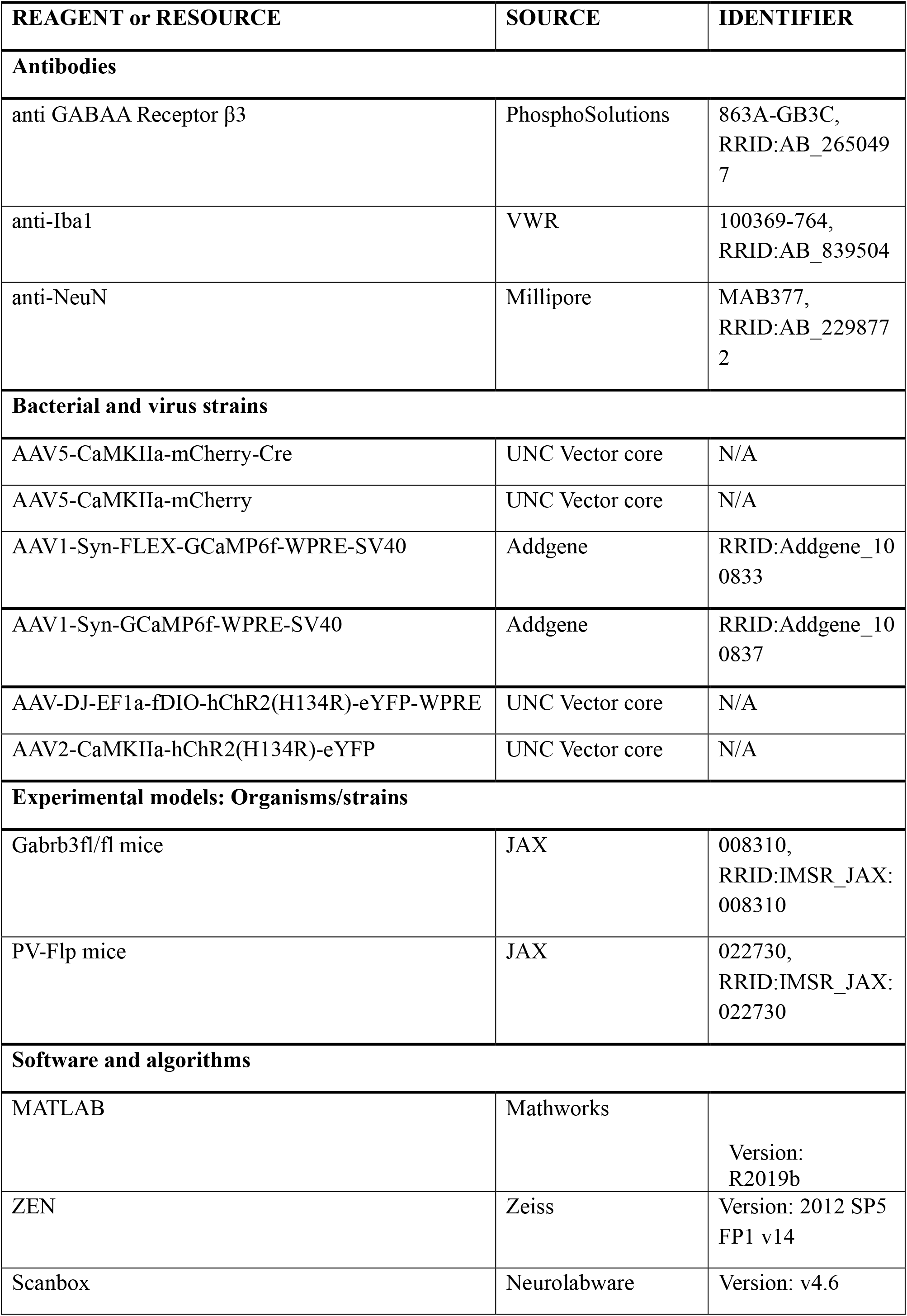
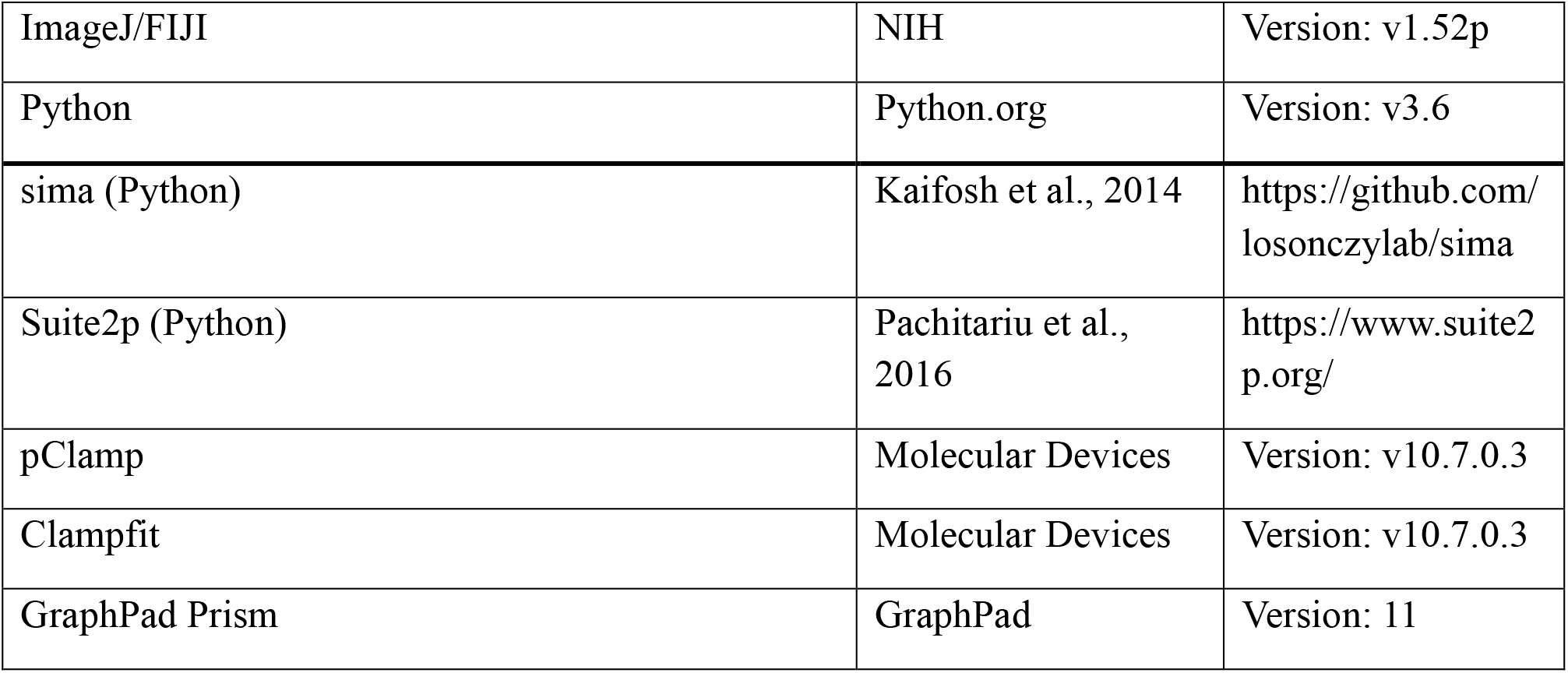

## STAR Methods

*Mice*: Gabrb3fl/fl mice (Strain #008310) and PV-Flp mice (Strain #022730) were obtained from Jackson Labs. All mice used were more than 7 weeks old and both male and female. Procedures were carried out in accordance with the National Institutes of Health guidelines for animal care and use and were approved by the Administrative Panel on Laboratory Animal Care of Stanford University (Protocol #30183). Mice were group housed (2–5 per cage), received ad libitum access to food and water, and were maintained on a 12 hr light/dark cycle throughout the study under standard housing conditions (21 ± 2 °C; 50 ± 15% humidity).

*Surgeries*: All surgeries were conducted under aseptic conditions using a small-animal stereotaxic instrument (AngleTwo, Leica Biosystems Inc.), UMP3 ultramicropump (WPI), and WPI syringes (Nanofil 33G Beveled). Mice were anaesthetized with isoflurane (5% for induction, 1.5–2.0% after) in the stereotaxic frame for the entire surgery and their body temperature was maintained using a heating pad. For electrophysiological recordings, Gabrb3fl/fl mice were stereotaxically injected with 250-300nL of AAV-CaMKIIa-mCherry-Cre or control AAV-CaMKII-mCherry virus (UNC vector core) into the right CA1 (AP: -2.0mm, ML: 1.4mm, DV: -1.45mm). For in-vivo electrophysiology, mice were implanted with a stainless steel headpost for head-fixation on a spherical floating ball as previously described^31,49^. For CA2 targeting, Gabrb3fl/fl mice were additionally injected with 200nL of AAV-CaMKIIa-hChR2(H134R)-eYFP (Addgene) virus into lateral CA2 (AP: -1.8mm, ML: 2.1mm, DV: -1.8mm), which achieved selective CA2 expression similar to recent reports of CA2 PCs being effectively transduced by AAV^50^. For 2p imaging, mice were injected with the same Cre or control AAVs as above bilaterally, and also injected with AAV-Syn-FLEX-GCamp6f or AAV-Syn-GCamp6f (Addgene). Mice were then chronically implanted with a chronic hippocampal imaging window and headpost for head-fixation on a linear treadmill as previously described^31,51,52^. To enable simultaneous electrophysiological recordings during 2p acquisition, a 4-channel silicon probe (Neuronexus, Q1x4-3mm-100-177-HQ4_21mm) was implanted contralateral to the imaging cannula. For ex-vivo electrophysiology Gabrb3 fl/fl mice were crossed to PV-Flp mice and resulting Gabrb3fl/fl;PV-Flp+ mice were injected with AAV-CamKIIa-mCherry-Cre and AAV-Ef1a-fDIO-ChR2-eYFP viruses (UNC vector core) into CA1.

*2p-imaging and ephys*: Mice were water restricted to 2mLs/day/mouse starting one week before and during 2p Ca2+ imaging. Head-fixed mice were imaged using a resonant scanner 2-photon microscope (Neurolabware), equipped with a pulsed IR laser (Mai Tai, Spectra-Physics), gated GaAsP PMT detectors (H11706P-40, Hamamatsu), and 16x objective (0.8 NA WI, Nikon). 2-photon image acquisition was controlled by a Scanbox (Neurolabware) system, which also synchronized behavioral video acquisition (Mako, Allied Vision), treadmill speed monitoring, and field potential recording via a DAC (National Instruments) to the imaging frames. Imaging of the pyramidal layer of area CA1 of the hippocampus was performed. Baseline signal 2-photon and hippocampal LFP were recorded while head-fixed mice were allowed to spontaneously run or rest on a cued linear treadmill. Data was collected at 15 fps with the laser tuned to 920 nm. LFP data was collected with open-ephys and synchronized with TTL pulses aligned to each acquired frame. The channel with the strongest signal for ripples and IEDs was selected for data analysis. Individual ripples/IEDs were initially screened based on a defined threshold (3 times the median absolute deviation of the signal) and manually confirmed. Image processing was performed as previously described^51,52^. To detect hypersychronous events, all traces were first smoothed using an exponentially weighted moving average and z-scored. For each frame, we counted the fraction of cells with zscore above 1. Then, we determined the average value of percent active cells in frames when the mouse was engaged in locomotion (speed > 0cm/s) to determine a threshold. Times at which the percent active cells exceeded this threshold during immobility and reached a peak were considered hypersynchronous events. We reported the frequency of these events for each mice and the percent active cells for each event. For determining recruitment of deep vs. superficial CA1PCs during hypersychronous events, we utilized these timepoints and reported the zscore at the peak of each event for each cell. Next, we divided the CA1PC layer along the midpoint between the stratum oriens and stratum radiatum, classifying cells falling on the oriens side as deep CA1 PCs and cells falling on the radiatum side as superficial PCs. Mean zscores were averaged across these anatomical distinctions and across all events in the recording. For calcium recordings where LFP signals could reliably record with IEDs (a subset of recordings were not reliably targeted to the CA1 PC layer), we aligned the frame at which the hypersynchronous event occurred to the LFP timepoint of when that frame was acquired. To further refine this timepoint for adequate LFP alignment, we averaged the calcium traces across all cells without smoothing.

*In-vivo LFP recording*: Bilateral craniotomies were performed over injection site and contralateral to it (same coordinates) under brief isoflurane anesthesia. Mice were then head-fixed on a floating spherical ball for free movement. Locomotor speed was recorded using synchronized video that recorded mouse behavior and ball movements, as previously described^49^. Two linear probes (A1x32-6mm-50-177-32, NeuroNexus) were slowly inserted into each hippocampal hemisphere until the deepest channels were approximately in the hilus. In a subset of recordings, the ipsilateral probe was a four-shank A1x32-Buzsaki32-CM32 (Neuronexus) to span multiple sites along the CA1 PC layer. Then, 30-60min recordings were obtained from mice as they behaved on a floating spherical treadmill using an Open Ephys acquisition system. For optogenetic stimulation experiments, only mice with verified CA2-specific expression of ChR2-eYFP were included (note: a small subset of mice had expression in CA3, which failed to evoke IEDs). A A1x32-Poly3-10mm-50-177-OCM32LP (Neuronexus) probe with a optical fiber mounted immediately dorsal to the most superficial channel was used. A 435nm DPSS-laser (Shanghai Laser & Optics Century) was connected to the probe to deliver light to the CA1, activating CA2 axons with 0.1msec to 100msec light pulses not exceeding 5mW measures at the fiber tip. Evoked response we measured using Data was analyzed using custom Python scripts. For spike detection, LFP recorded was first bandpass filtered between 3 and 50 Hz, and the oriens channel was subtracted from the radiatum channel to result in a differential recording. Differential signals exceeding 0.5mV were detected as spikes as long as they were more than 100msec apart from other detected spikes. For ripple/HFO detection, LFP from the CA1 PC layer was bandpass filtered between 100 and 600 Hz and a Hilbert transformation was performed to derive the envelope and instantaneous frequency. Ripple/HFOs were included if the envelope reached 5 standard deviations, exceeded 3 standard deviations for 15 msec, and were more than 125msec apart. The ripple frequency for each event was averaged across 8 msec centered on the peak of the envelope, which was used to classify the event as a ripple (<200Hz) or HFO (>200Hz). For radiatum amplitude during each detected event, the maximal absolute voltage within 10msec on either side of the envelop peak was taken. Ripples and HFOs were considered bilaterally synchronous if they well within 100msec of each other.

*Ex-vivo electrophysiology*: At 2–5 weeks following initial injection, mice were deeply anesthetized by Ketamine/Xylazine and then transcardially perfused with an ice-cold protective recovery solution containing (in mM): 92 NMDG, 26 NaHCO3, 25 glucose, 20 HEPES, 10 MgSO4, 5 Na-ascorbate, 3 Na-pyruvate, 2.5 KCl, 2 thiourea, 1.25 NaH2PO4, 0.5 CaCl2, titrated to a pH of 7.3–7.4 with HCl70. Coronal slices (300 µm) containing the hippocampus were cut in ice-cold protective recovery solution using a vibratome (VT1200S, Leica Biosystems). Brain slices were then incubated in 35 °C protective recovery solution for 12 min. Subsequently, brain slices were maintained in room temperature aCSF consisting of (in mM): 126 NaCl, 26 NaHCO3, 10 glucose, 2.5 KCl, 2 MgCl2, 2 CaCl2, 1.25 NaH2PO4. All solutions were equilibrated with 95% O2/5% CO2.

Intracellular recordings were performed in a submerged chamber perfused with oxygenated aCSF at 3 ml/min and maintained at 33 °C by a chamber heater (BadController V, Luigs and Neumann). CA1 neurons were visualized using DIC illumination on an Olympus BX61WI microscope (Olympus Microscopy) with an sCMOS camera (Flash 4.0 LT+, Hamamatsu). Epifluorescence illumination from an LED lamp (Solis-3C, Thorlabs) was used to identify transfected neurons and deliver optogenetic stimulation pulses (1ms, 0.1Hz). Recording pipettes were pulled from thin-walled borosilicate capillary glass (King Precision Glass) using a P97 puller (Sutter Instruments) and were filled with (in mM): 126 K-gluconate, 10 HEPES, 4 KCl, 4 ATP-Mg, 0.3 GTP-Na, 10 phosphocreatine (pH-adjusted to 7.3 with KOH, osmolarity 290 mOsm), as well as 0.2% biocytin. Pipettes had a 3–5 MΩ tip resistance.

Whole-cell recordings were performed on mCherry-positive CA1 neurons in the dorsal hippocampus (A/P: −1.5 to 2.4 mm). Pipette capacitance was neutralized for all recordings and cells were held at a potential of −65 mV. Deep cells were targeted as the first or second cell in the pyramidal layer immediately adjacent to oriens. Superficial cells were targeted as the first or second cell in the pyramidal layer immediately adjacent to radiatum. Data were acquired in pClamp software (Molecular Devices) using a Multiclamp 700B amplifier (Molecular Devices), low-pass filtered at 2 kHz, and digitized at 10 kHz (Digidata 1440A, Molecular Devices). Data analysis was performed using Clampfit.

*Immunohistochemistry*: Mice were deeply anesthetized with a mixture of ketamine and xylazine (80–100 mg/kg ketamine, 5–10 mg/kg xylazine; intraperitoneal) and transcardially perfused with 10 mL of 0.9% sodium chloride solution followed by 40 ml of cold 4% PFA dissolved in phosphate buffer solution. The excised brains were held in a 4% PFA solution for at least 24 h before being sectioned into 60 μm slices using a vibratome (Leica VT1200S; Leica Biosystems Inc.). After the initial TBS rinses, free-floating FC sections were incubated for 1 h at room temperature (RT) in TBS+ (1× TBS with 1% BSA and 0.05% Triton X-100) to block non-specific binding. Sections were then incubated for at least 17 h at RT on a gentle rocker in TBS+ containing primary antibody solution (rabbit anti-Iba1 1:1000; mouse anti-NeuN 1:500; rabbit anti GABAA Receptor β3 1:300).

Following primary incubation, sections were washed in 1× TBS (4 × 10 min) and incubated for 2 h at RT with secondary antibody (goat anti-rabbit Alexa488 1:1000; goat anti-mouse Alexa405 1:1000). Sections were then washed in 1× TBS (4 × 10 min), mounted onto Superfrost Plus charged slides, air-dried, and coverslipped with VECTASHIELD Antifade Mounting Medium (Vector Laboratories). Imaging was performed on a Zeiss LSM 800 confocal microscope using a 20x objective. For each animal, 3–5 regions of interest were imaged for each region corresponding to CA1 and a z-stack of 5–7 images was taken. Imaging conditions were conserved across all regions and sections imaged.

